# Piezo3 is a novel mechanosensitive Piezo ion channel in vertebrates

**DOI:** 10.64898/2026.05.15.725496

**Authors:** Ziyu Dong, Dingxun Wang, Beichen Wang, Jillian A. New, Yuk Fai Leung, GuangJun Zhang

**Author notes:** Corresponding author: GuangJun Zhang. These authors contributed equally to this work (Ziyu Dong, Dingxun Wang).

## Abstract

Mechanosensing and mechanotransduction are essential for all living cells. In mammals, Piezo1 and Piezo2 are two mechanically activated cation channels that serve as mechanosensors for a variety of physiological and pathological processes, ranging from touch sensing to sickle cell disease. These two channels are well evolutionarily conserved, and orthologous genes can be traced back to the origin of vertebrates, which underwent whole-genome duplications (WGDs). The number of paralogous genes originating from the vertebrate WGD varies across gene families. Thus, whether there are more *PIEZO* paralogous genes in vertebrates remains understudied. Here, we identified *piezo3*, a new paralog of the piezo gene family, and analyzed its evolutionary history using phylogenetic and synteny analyses. The *piezo3* gene is present in most vertebrate lineages but absent in birds and most mammals, likely due to nonfunctionalization after WGDs. In addition, we demonstrated that this channel could mediate calcium flux in response to mechanical stimuli in HEK293T cells, suggesting that Piezo3 exhibits PIEZO1/2-like activation and conduction channel functions. Our CRISPR mutation analysis revealed that the zebrafish *piezo3* gene is not developmentally essential, possibly because its expression overlaps with other PIEZO channels. Mutant zebrafish showed elevated sensitivity to mechanical force and increased locomotor activity under (photopic) light illumination. Our results suggest that this new mechanical-sensing Piezo channel is widespread in vertebrates and may be critical for vertebrate adaptation by modulating mechanical sensing and light responses during evolution.

**SIGNIFICANCE:** All living cells must sense mechanical forces, whether endogenous or exogenous, and respond to them by transforming these forces into biological signals, which is essential to a wide range of cellular processes, including cell division, growth, and differentiation. PIEZO channels are well-characterized, critical, versatile mechanotransducers for touch and pain physiology and for human diseases. Currently, PIEZO1 and PIEZO2 are the only two known PIEZO channels in most vertebrates. In zebrafish, there are two Piezo2 channels (Piezo2a and Piezo2b) due to extra genome duplication in the ray-finned fishes. Here, we report Piezo3 channel, a long-missing paralog of Piezo1 and Piezo2, in most vertebrates. This channel is present in the majority of vertebrate lineages, except for most birds and mammals. The zebrafish *piezo3* gene is expressed during early embryogenesis, and mutation of this gene leads to zebrafish larvae responding to tapping mechanical force and light with active movement. The widespread distribution of this Piezo3 channel across most vertebrate species, but its absence in birds and most mammals, suggests it may play important roles in vertebrate physiology and evolution.

## INTRODUCTION

Mechanosensing and mechanotransduction are fundamental cellular processes that are manifested at many levels of cellular behaviors, including cell division, growth, and the transmission of biochemical signals (1). Ion channels play an important role in transducing mechanical force into electrochemical responses by allowing ions to cross cell membranes along their concentration gradient. MscL (large conductance mechanosensitive ion channels) was the first mechano-gated channel identified as an osmotic stress-release valve in bacteria (2). Homologous MscL was reported in Archaea but not in mammals (1). It took decades of searching before the PIEZO1 and PIEZO2 channels were identified (3). After that, the orthologs of these two genes were extensively investigated in various organisms, especially the mouse (4, 5). Piezo channel genes are not limited to vertebrates; invertebrates and plants possess piezo channels (6-9). Now, the mechanotransduction by PIEZO1 and PIEZO2 channels is increasingly being incorporated into many biological research fields (10). PIEZO1 is broadly expressed across many tissues and organs and plays critical roles in cardiovascular, skeletal, and digestive physiology. PIEZO2 is mainly expressed in the ganglion neurons and is responsible for somatosensory, auditory, respiratory, and urinary sensing (11, 12).

Whole-genome duplications (WGDs) play a significant role in shaping paralogous gene numbers and expanding developmental genetic toolkits that underlie vertebrate morphological novelties and diversity (17-19). It is well accepted that there are two rounds of WGDs (1R and 2R) around the origin of vertebrates (17-20). All the genes of vertebrate ancestors were duplicated twice, creating four ohnologs through the two rounds of WGDs. One or more of these four ohnologs (paralogs generated by WGD) may be lost in vertebrate lineages. For example, there is one fibrillar collagen and one Hox gene cluster in invertebrates, while there are four fibrillar collagen Hox gene clusters in tetrapods (21). After the vertebrate lineage split, another round of WGD (3R) occurred in the common ancestor of all extant teleosts (22). This 3R resulted in more ohnologs in teleost species, although many duplicated genes were lost (23, 24). For example, there are multiple potassium channel duplicates, *i*.*e*., ohnologs, in zebrafish (16, 25, 26). These duplicated genes play an essential role in embryogenesis and the diversity of vertebrate morphological structures, such as skeletons, through subfunctionalization, neofunctionalization, and gene loss (18, 20, 21, 27, 28). PIEZO1 and PIEZO2 have been extensively investigated for mechanotransduction in mice and other species, while the evolutionary history of this type of channels in vertebrates remains further investigated, especially in teleosts, given the three rounds of WGD events in vertebrates.

Zebrafish is one of the most common vertebrate model systems for current biomedical research due to numerous advantages, such as external embryonic development, early transparent embryos, tractable genetics, and vertebrate biology that is homologous to humans (29, 30). Moreover, its phylogenetic position and its genome, which underwent 3R, made the zebrafish model attractive for evolutionary functional studies. Here, we report a new vertebrate *piezo* gene, *piezo3*, with its evolutionary history. We found that the *piezo3* gene is present in most vertebrate lineages but is lost in birds and most mammals. In addition, we examined its expression patterns, which suggest Piezo3 may play a role in embryogenesis and physiology. Our CRISPR mutants support this notion. Although the piezo3 gene is not developmentally essential, it modulates zebrafish larval responses to mechanical and photopic stimuli.

## RESULTS

### Identifying a new piezo channel gene in zebrafish

The two-pore domain potassium (K2P) channels are also involved in mechanosensing and transduction (13-15). Recently, we revealed that vertebrate K2P potassium channels were expanded by WGDs (16). It is reasonable to expect that WGDs also shape PIEZO channels, and we hypothesized that more paralogous genes of *piezo1* and *piezo2* may exist in teleosts. To test this hypothesis, we chose zebrafish because the functions of *piezo* genes (*piezo1, piezo2a*, and *piezo2b*) have already been reported, and the zebrafish genome is relatively well-annotated (31, 32). First, we searched for *piezo* genes in the zebrafish genome (GRCz11) using already-known zebrafish *piezo1* as bait by BLAT. We identified *si:dkey-11f4*.*7* as a new paralog of the *piezo1* gene. In ZFIN, there are six transcripts from this gene (*si:key-11f4*.*7-201* to *si:key-11f4*.*7-206*) that correspond to six transcripts of the gene ENSDARG00000099207 in ENSEMBL. The sizes of these six transcripts (443-972 bps) are much shorter than the *piezo1* transcript. In contrast, in NCBI, the *si:dkey-11f4*.*7* gene (Gene ID: 569687) has two much longer transcripts (XM_068223521.1 and XM_009304390.4), and they code two putative proteins (2605aa and 2651aa). This discrepancy likely resulted from the incomplete zebrafish genome annotation. The *si:key-11f4*.*7* shared homology with the current three known zebrafish *piezo1, piezo2a*, and *piezo2b* (**Table 1**). It is also worth noting that the *piezo2a* gene is annotated as two genes (*piezo2*.*a* and *piezo2a*.*1*) in ZFIN, and both are partial sequences of piezo2a (**Table 1**).

**Table 1.**
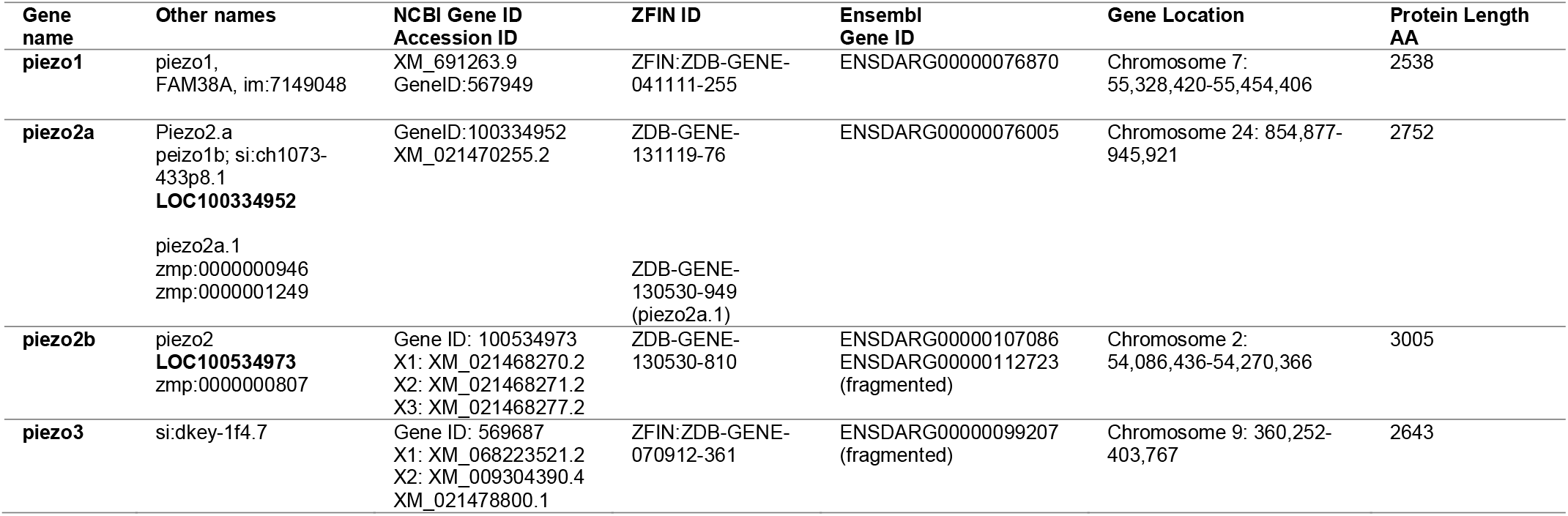
Zebrafish *piezo* paralogous genes.

### Vertebrate *piezo* genes may have been duplicated through WGDs

This new *si:dkey-11f4*.*7* gene could be unique in zebrafish or conserved in other vertebrates. To address this question, we used this gene as bait and searched NCBI and Ensembl for homologous sequences. This gene is also present in most vertebrate lineages, including chondrichthyans, holostei, teleosts, amphibians, and reptiles, but not in currently sequenced birds and most mammals. To further clarify its relationships with PIEZO1 and PIEZO2, we conducted phylogenetic analyses using maximum likelihood and Bayesian methods. Both analyses yield consistent results. In vertebrates, there are three clusters: PIEZO1, PIEZO2, and Si:dkey-11f4.7. The Si:dkey-11f4.7 clade is closer to the PIEZO2 than the PIEZO1 clade (**Fig. 1** and **Fig. S1**). Thus, we suggest renaming the *si:dkey-11f4*.*7* as PIEZO3 based on our phylogenetic analysis. Most teleost species possess duplicated *piezo2* genes (*piezo2a* and *piezo2b*), indicating they originated from the third round WGD (3R). Interestingly, the PIEZO3 gene is present in some mammalian genomes, such as those of the armadillo, red deer, and tree shrew, suggesting that the loss of this gene might not have occurred in the ancestor of mammals, but rather independently in each sublineage.

**Figure 1.**
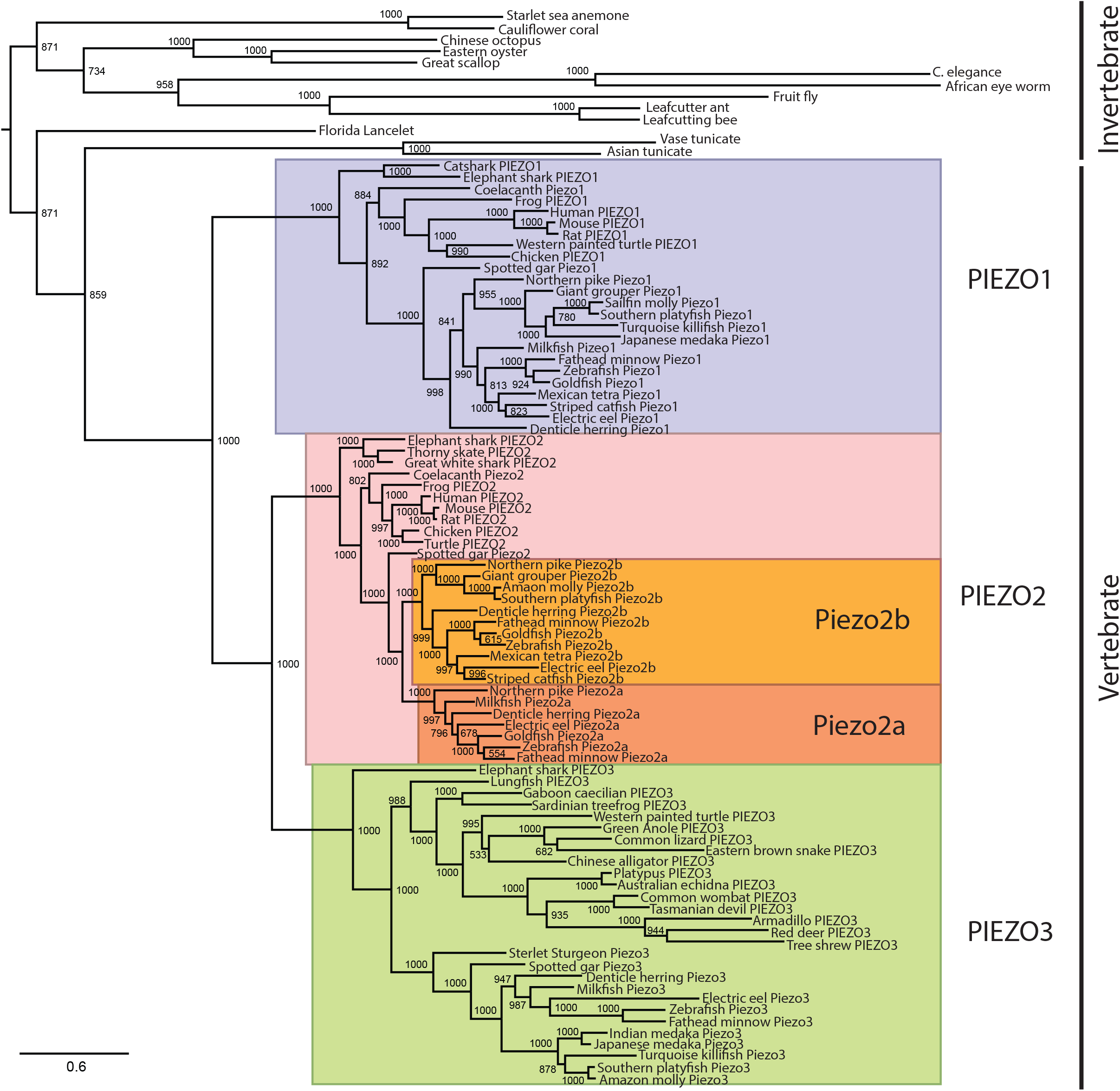
Maximum likelihood phylogeny of animal PIEZO channels. Some representative species (human, mouse, zebrafish, spotted gar, shark, lancelet, tunicate, fruit fly, and worm) were analyzed. Numbers at each node indicate the supporting values based on 1,000 bootstrap replicates. Branch lengths are proportional to the expected number of replacements per site. The phylogeny was inferred using the JTT model with a gamma distribution in PhyML. All vertebrate PIEZOs were grouped together, and the three types of PIEZO channels formed a distinct clade, which is color-coded and indicated with vertical black lines. The teleost Piezo2 channels were also color-highlighted. All invertebrate PIEZO channels formed a distinct outgroup on the phylogenetic tree.

To further confirm whether the vertebrate piezo genes are true paralogs generated by WGDs, we conducted syntenic analyses across a few representative species. We found that *piezo1* is located in an evolutionarily conserved synteny (*aprt-cdt1-piezo1-ctu2-rnf166-snai3*), and *piezo2* is linked with *napg* and *apcdd1* genes (**Fig. 2A-B**). Similarly, piezo3 is located in a longer synteny (*prelid3b-atp5f1e-tubb1-ctsz-nelfcd-gnas-piezo3-npepl1-stx6-apcdd1l-vapb-rab22a*). This *piezo3* synteny becomes shorter and more variable in teleosts (Fig. 2C). The linkages of piezo2-apcdd1 and *piezo3-apcdd1l* are consistent with their closer clustering in our phylogeny analysis, suggesting they may have arisen from a common ancestor during the teleost-specific WGD (3R).

**Figure 2.**
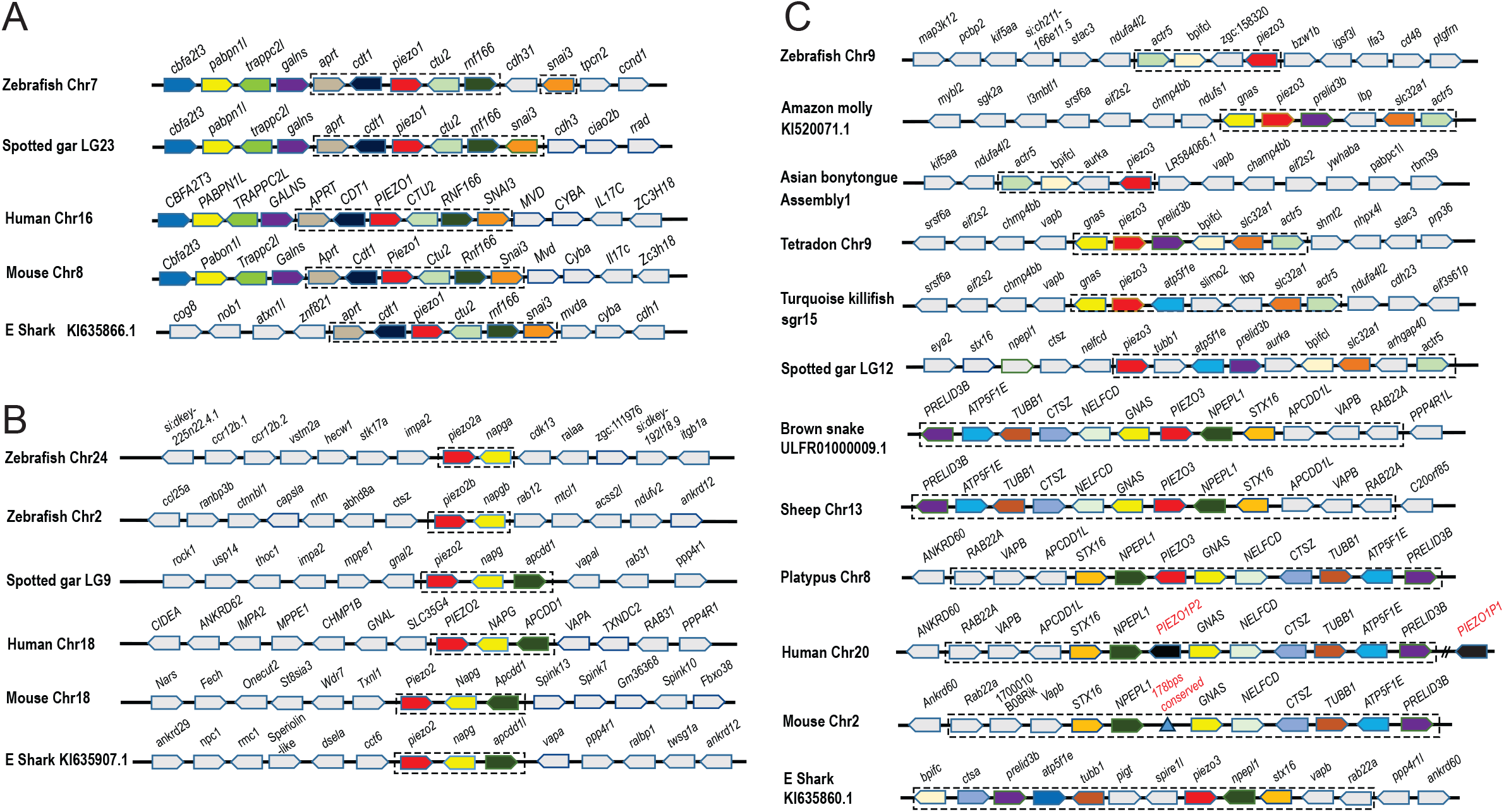
Syntenic analyses of the piezo genes across representative vertebrate species. The Illustration of the genes and their sizes is not proportional to the actual distances between genes. **A**. Common vertebrate piezo1 synteny. The *piezo1* gene was highlighted in red. The synteny (*aprt-cdt1-piezo1-ctu2-rnf166*) is shared among all the representative species. **B**. Common vertebrate piezo2 synteny (*piezo2-napg-apcdd1*). The *piezo2* is highlighted in red. The synteny was boxed with blue dashed lines. After duplication, the *apcdd1* gene is lost in both *the piezo2a and piezo2b synteny blocks*. **C**. Common vertebrate *piezo3* synteny. A few more representative species were included for comparison. The *piezo3* synteny is boxed with dashed lines. The gene order of this synteny is conserved among sharks, snakes, and mammals, but slightly shifted in spotted gar and teleost. The *PIEZO3* genes become pseudogenes (red letters) in humans and mice.

To reveal how the *PIEZO3* gene was lost in humans, we further analyzed the sequences of the conserved synteny. According to the current human genome (GRCh38.p14), there is a pseudogene, *PIEZO1P2*, between *NPEPL1* and *GNAS* (Chromosome 20: 58,740,532-58,783,821). *PIEZO1P2* encodes a 2161bp transcript and shares homology with *PIEZO1*, supporting that this processed *PIEZO1P2* pseudogene transcript resulted from *PIEZO3* inactivation during Evolution. Interestingly, at the downstream of this synteny, there is another pseudogene, *PIEZO1P1* (219bp), which is likely a rudimentary part of *PIEZO3* (**Fig. 2C**). Similarly, we found a 178bp conserved region (chromosome 2:174075082:174075795) in the mouse genome (GRCm39). It has 87.72% identity to human PIEZO1P2, indicating that the mouse Piezo3 gene was also inactivated/lost in the same way. According to the GTEX portal, *PIEZO1P2* is expressed in the spleen and testis, although its function remains unknown (**Fig. S2**).

### Zebrafish *piezo* genes’ expression during embryogenesis

As PIEZO3 is a new vertebrate piezo channel, no functional study has been conducted. To gain insight into its functions and validate that *piezo3* is an actively expressed gene, we performed whole-mount *in situ* hybridization during zebrafish embryogenesis. This gene was not turned on until 5-somite stage, and it is mainly expressed in neuronal tissues (**Fig.3 A-F**). At the 15-somite stage, it was expressed at the ventral side of the brain (**Fig. 3G-I**). From the 21-somite to 24hpf stage, *piezo3* was mainly expressed in the brain and the eye, including the lens (**Fig. 3J-M**). The brain and eye expression remained but became less expressed at 48 hpf (hours post fertilization). Evident expression was detected in pharyngeal arches and pectoral fin buds (**Fig. 3N-O**). This expression of the pharyngeal arch and pectoral fin buds continued to the 72hpf stage. In contrast, the head expression was reduced (**Fig. 3P-Q**), suggesting Piezo3 might play a role in neural, pharyngeal, and pectoral fin development.

**Figure 3.**
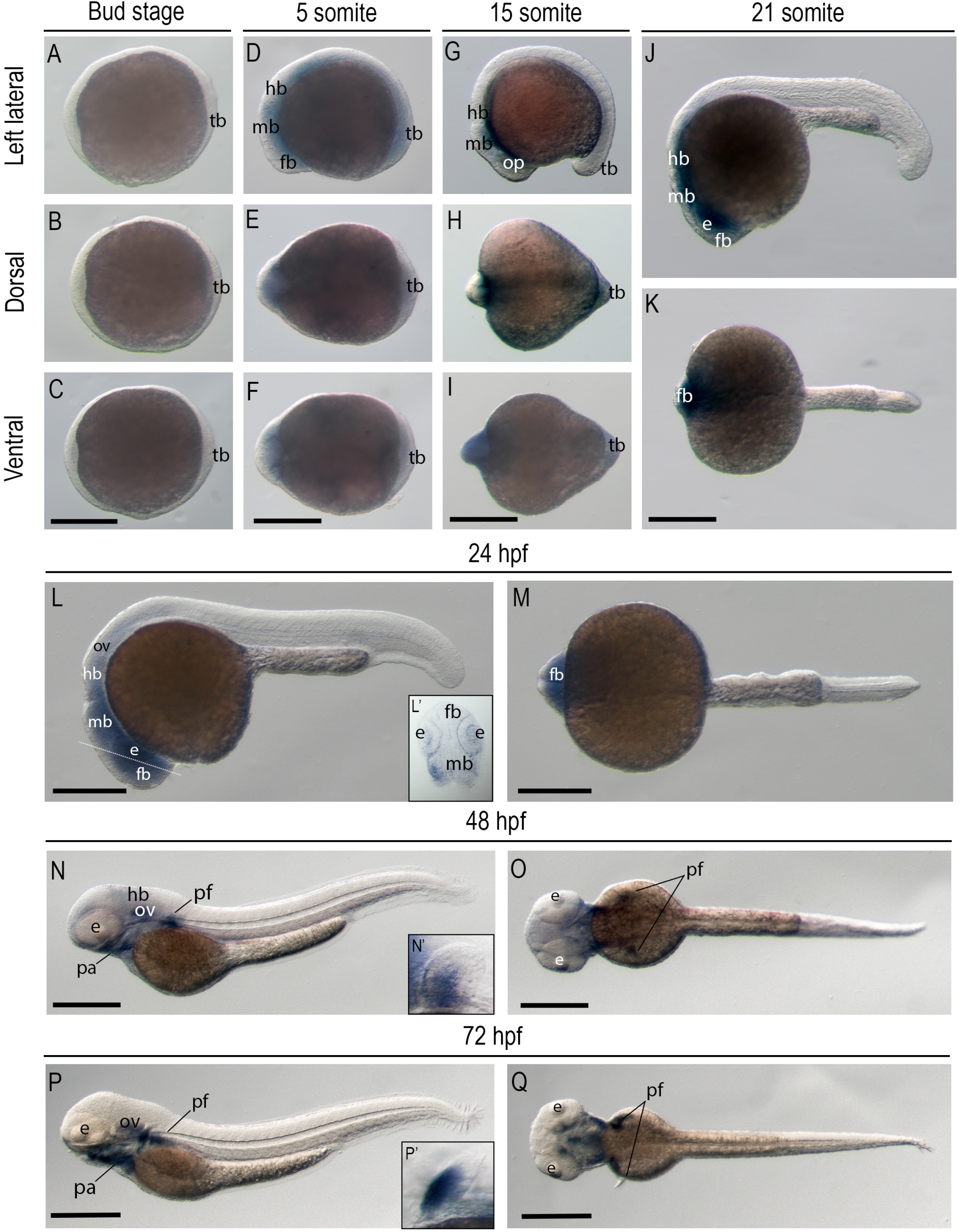
Zebrafish *piezo3* gene expression during zebrafish embryogenesis. Whole-mount in situ hybridization of zebrafish embryos at bud stage (**A**-**C**), 5 somite stage (**D**-**F**), 15 somite stage (**G**-**I**), 21 somite stage (**J**-**K**), 24hpf (**L**-**M**), 48hpf (**N**-**O**), and 72 hpf (**P**-**Q**). The anterior is to the left in all the whole-mount images, and the dorsal is to the top in all transverse sections (**L, N, P**). The piezo3 gene expression is shown laterally (**A, D, G, J, L, N, P**). Dorsal view of gene expression of *piezo3* (**B, E, H, K, M, O, Q**). Ventral view of gene expression of *piezo3* (**C, F, I**). The white dashed lines indicate the proximate positions of the sections shown in the insert (**L’**) of the same panel (**L**). Magnified pectoral fin buds are highlighted in the inserts (**N’, P’**) in the corresponding panels (**N, P**). Scale bars were added to the left of the bottom row. 250[μm for whole-mount images. *e*, eye; *fb*, forebrain; *hb*, hindbrain; *mb*, midbrain, *op*, optic vesicles; *ov*, otic vesicles; *pa*, pharyngeal arches; *pf*, pectoral fins; *tb*, tail bud.

The *piezo3* gene is paralogous to *piezo1, piezo2a*, and *piezo2b*. To elucidate whether they are coexpressed during zebrafish embryogenesis, we also systematically analyzed the three paralogs. The *piezo1a* was first detected in the forebrain during the late gastrulation, the bud stage, and continued to the 5-somite stage (**Fig. S3A-F**). From the 15-somite to the 21-somite stage, it was also expressed at the otic vesicles and ventral somite mesoderm (**Fig. S34G-K**). This ventral somite and otic vesicle expression continued to 24hpf (**Fig. S3L-M**). This expression is consistent with its functions in red blood cells and the vertebral column (32, 33). At 48hpf, the somite expression retracted, but the *piezo1* transcript was detected at the otic vesicle, pharyngeal arch, and pectoral fin buds. These expression domains remained at 72hpf (**Fig. S3P-Q**).

The *piezo2a* gene was first detected in the forebrain at the 5-somite stage and was lightly expressed in the neural tube (**Fig. S4A-F**). The forebrain domain continued at the 15-somite stage (**Fig. S4G-I**). From the 21-somite to the 24hpf, *piezo2a* was mainly expressed at the ventral brain and ventral mesoderm of somites (**Fig. S4J-M**). Similar to *piezo1*, this domain may be related to vertebral column development (32). At 48hpf, the ventral mesoderm recessed, while the pharyngeal arches, pectoral, and caudal fin buds started to express the *piezo2a* gene (**Fig. S4N-O**). These expression patterns persisted to 72 hpf and 5 dpf (days post-fertilization) (**Fig. S4P-S**). Interestingly, its expression within caudal fin buds is relatively stable at the proximal part. While *piezo2a* within pectoral fin buds is dynamic: most mesenchyme (48hpf), basal mesenchyme (72hpf), and the interface between epithelial and mesenchyme (5dpf), suggesting this gene might have an essential role in fin development. The *piezo2b* gene is expressed mainly in Rohon-Beard neurons of the neural tube and the trigeminal ganglia (**Fig. S5A-Q**), consistent with a previous report (31).

### Zebrafish Piezo3 is a calcium-conducting cation channel in HEK293T cells

PIEZO1 and PIEZO2 selectively conduct Ca^2+^, Na^+^, K^+^, and Mg^2+^, and both are sensitive to streptomycin (3). To examine whether zebrafish Piezo3 has a similar function, we first took advantage of AlphaFold to examine its 3D structure using the available amino acid sequences. Indeed, the zebrafish Piezo3 has a very similar 3D structure to human PIEZO1, PIEZO2, and the other three zebrafish Piezo1, Piezo2a, and Piezo2b (**Fig.S6 A-E**). Next, we cloned the full-length zebrafish *piezo3* into a CMV promoter-driven plasmid construct with an IRES (Internal Ribosome Entry Site)-mediated EGFP marker. We co-transfected it with a Ca^2+^ reporter, jRGECO1a, into HEK293T cells. After transfection, there are four types of cells: 1) Cells with both EGFP and red fluorescence (Piezo3 and jRGECO1a), green-only (Piezo3), red-only (jRGECO1a), and no fluorescence cells (un-transfected). To study the effects of Piezo3, we compared the red fluorescence intensity (Calcium readout) between red-only cells (no Piezo3) and both green and red cells (Piezo3 and jRGECO1a) (**Fig. 4A-B**). We observed a slightly higher baseline fluctuation in the Piezo3 expression cells (**Fig. 4C, H**). To test their mechanical response, we applied a gentle force using a blunt-end glass probe mounted on a micromanipulator. A rapid rise in red signal transient and a slower signaling decay were observed (**Fig. 4D, H**), indicating calcium ions were moving through the Piezo3 channel. To test whether the Piezo3 is sensitive to streptomycin, we added streptomycin at two concentrations to the cell medium and repeated the stimulation experiments. We found that the cells showed significantly smaller or no changes in red signal (**Fig. 4E, H**). The Piezo3-expressing cells responded to mechanical force (**Fig. 4F, H**) at a low streptomycin concentration. However, the cells failed to respond to probe poking at the higher streptomycin concentration (**Fig. 4G-H**). Overall, these results suggest that zebrafish Piezo3 is a functional mechanical channel, similar to Piezo1 and Piezo2.

**Figure 4.**
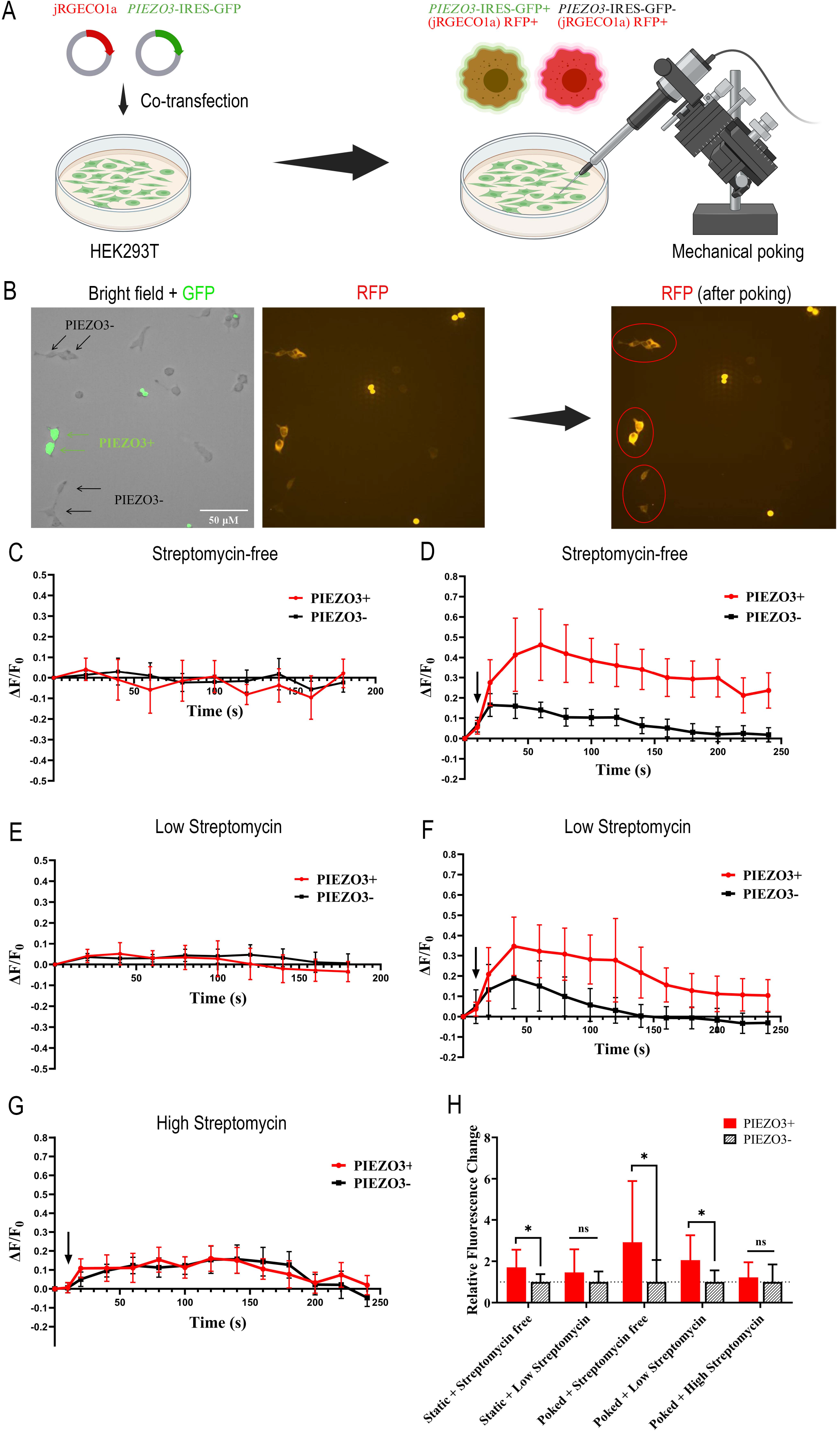
Piezo3 mediated intracellular calcium signals in response to mechanical force in HEK293T cells. **A**. Experimental design for co-transfection and calcium signal measurement of the mechanical force-induced response. **B**. Representative images of the RFP+/GFP+ and RFP+/GFP-HEK293T cells in untouched or mechanically stimulated conditions. The merged image of the bright-field and GFP channels showed piezo3-expressing GFP+ and GFP-cells, and the RFP channel showed jRGECO1a-expressing RFP+ cells. The red rectangle labeled the cells that were mechanically stimulated. **C**-**D**. Calcium signal responses of the RFP+/GFP+ and RFP+/GFP-HEK293T cells in streptomycin-free culture conditions without (**C**) and with (**D**) glass probe touch. **E**-**F**. Calcium signal responses at a low streptomycin (172μM) condition without (**E**) and with (**F**) glass probe touch. **G**. Calcium signal responses at a high streptomycin (1032μM) condition with glass probe touch. The vertical black arrow indicates the time of glass probe application. RFP was recorded at 0.5 frames/s for 3-4 min. ΔF/F_0_ (ΔF=F-F_0_) was used to describe the calcium signal response (mean + 95%CI). 10-15 cells were recorded for each group. **H**. Comparisons of the overall calcium signal responses between PIEZO3-expressing and non-PIEZO3 HEK293T cells. The total area under the curve (AUC) over the ΔF/F_0_=0 baseline was calculated to quantify the overall calcium signal response for the record duration. *p<0.05.

### Zebrafish piezo3 is developmentally nonessential and may be responsible for larval mobilization

To investigate the biological functions of the zebrafish Piezo3 channel, we mutated this gene using CRISPR-Cas9 (34). Three loss-of-function mutants, *pu117* (large knockout, LKO), *pu118* (indel and small knockout, SKO), and *pu119* (indels) were generated and validated by sequencing (**Fig. 5A-B**). To eliminate potential off-target mutations, we outcrossed F1 mutants to WT fish in established fish lines. Then, homozygotes were generated by in-crossing F_2_ adults for phenotype assessment. Homozygotes are fertile and show no evident morphological alterations. Consistently, we observed an approximate Mendelian ratio with the three mutants (**Fig. 5C**), suggesting that the *piezo3* gene is developmentally nonessential. As *piezo3* was expressed in the eyes of 1dpf fish embryos (**Fig. 3L**), we first analyzed the vision of 7dpf *pu117* larval zebrafish using the VMR assay. We found that the mutant displacement distance was greater than that of the wildtype sibling control, although there was no difference in startling response (**Fig. 5D**). Moreover, the pu117 mutant is more sensitive to mechanical tapping compared to the wildtype sibling control (**Fig. 6E**), suggesting that Piezo3 might have a repressive function in larval fish mobilization. We did not detect histological differences in the adult retina (**Fig. 5F-I**). As piezo3 was also expressed in the pharyngeal arches (**Fig. 3N, P**), we next examined *pu117* larva and adult gill arches and found no cartilage changes (**Fig. 5J-U**). As piezo1 plays an essential role in skeletogenesis and is responsible for intramuscular bones in zebrafish and carp (35), we then examined the adult zebrafish skeletons using alizarin red staining. However, we did not find an evident abnormality of overall axial and intramuscular bones (**Fig. 5W-AC**).

**Figure 5.**
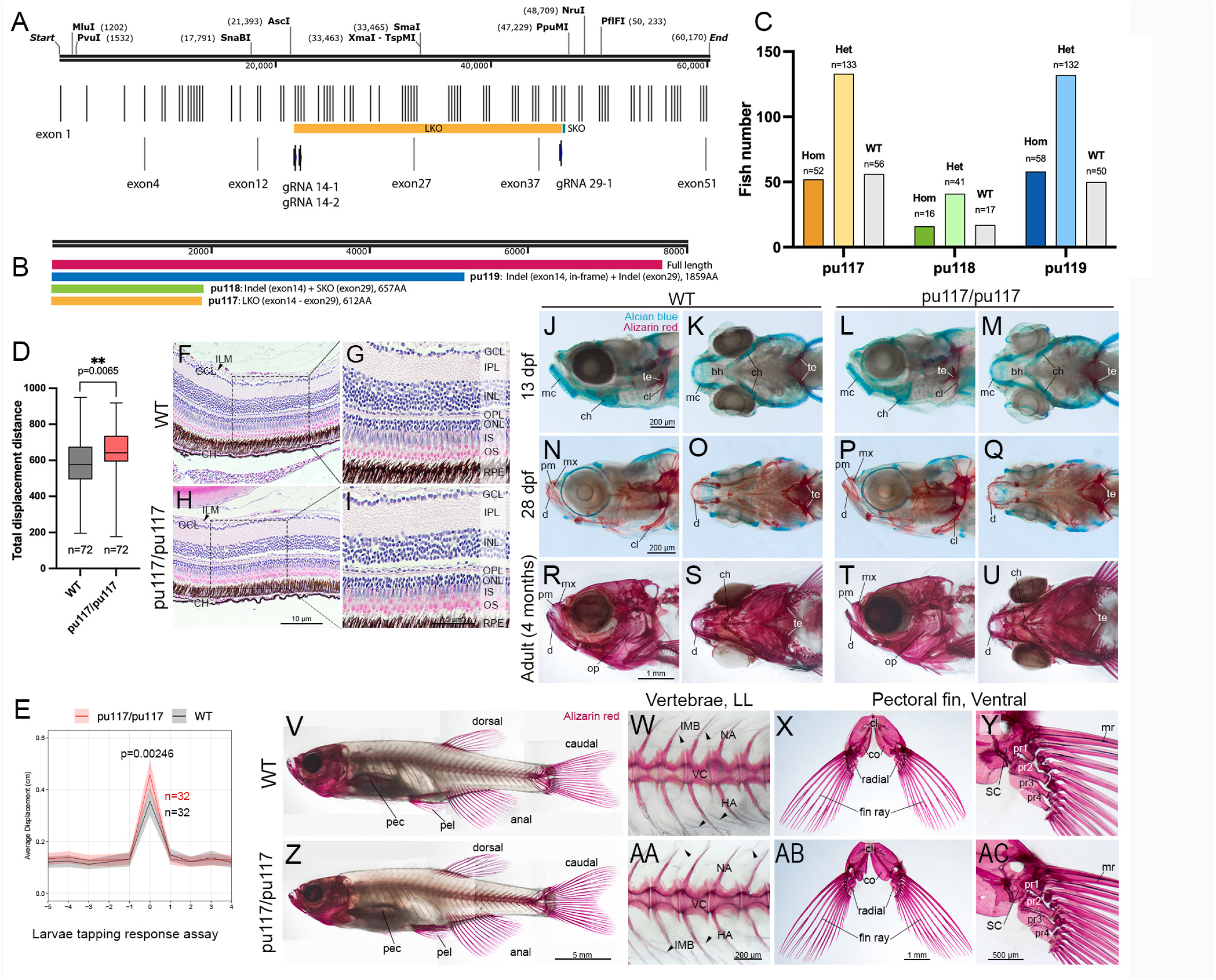
Zebrafish *piezo3* is a developmentally nonessential gene. **A**. Illustration of the *piezo3* locus based on the genomic fragment sequence, CR753874.9 in NCBI. Exons are shown as black vertical lines. Three CRISPR gRNA target sites are highlighted by blue vertical bars. The large knockout (LKO) truncated region is indicated by an orange bar, and the small knockout (SKO) region is shown as a green bar. **B**. Predicted *piezo3* proteins of full frame and 3 mutant alleles generated by CRISPR-Cas9: pu117 (LKO, exon 14 – exon 29), pu118 (exon 14 indel mutation + exon 29 SKO), pu119 (exon 14 in-frame indel mutation + exon 29 indel mutation). **C**. Genotype distribution from a F2 heterozygote in-crosses from three *piezo3* mutant lines (pu117, pu118, pu119). Genotypes are close to the expected Mendelian ratios. **D**. Larval displacement by mechanical tapping. **E**. Movement following mechanical stimulation by 3 independent visual motor response (VMR) assays. **F-I**. Hematoxylin and eosin (H&E) staining of adult zebrafish eyes in wild type (WT) and *piezo3* LKO homozygous fish (pu117/pu117). *CH*, choroid; *GCL*, ganglion cell layer; *ILM*, inner limiting membrane; *INL*, inner nuclear layer; *IPL*, inner plexiform layer; *IS*, inner segments; *ON*, optic nerve; *ONL*, outer nuclear layer; *OPL*, outer plexiform layer; *OS*, outer segments; *RPE*, retinal pigment epithelium. **J-U**. Alcian blue and Alizarin red staining of 13 dpf larvae **(J-M)**, 28 dpf juveniles **(N-Q)**, and adult **(R-U)** WT and *piezo3* LKO homozygous fish. *bh*, basihyal; *ch*, ceratohyal; *cl*, cleithrum; *d*, dentary; *LL*, left lateral; *mc*, Meckel’s cartilage; *mx*, maxilla; *pm*, premaxilla; *op*, opercle; *te*, teeth; *V*, ventral. **V-AC**. Alizarin red staining of the adult skeleton in WT and pu117 homozygous fish. Whole fish (**V, Z**) and higher magnification views of vertebrae (**W, AA**) and pectoral fin (**X, Y, AB, AC**) are shown. *anal*, anal fin; *caudal*, caudal fin; *cl*, cleithrum; *co*, coracoid; *dorsal*, dorsal fin; *HA*, haemal arches; *IBM*, intermuscular bone; *mr*, marginal ray; *NA*, neural arches; *pec*, pectoral fin; *pel*, pelvic fin; *pr1-4*, proximal radials; *SC*, scapula; *VC*, vertebral column.

**Figure 6.**
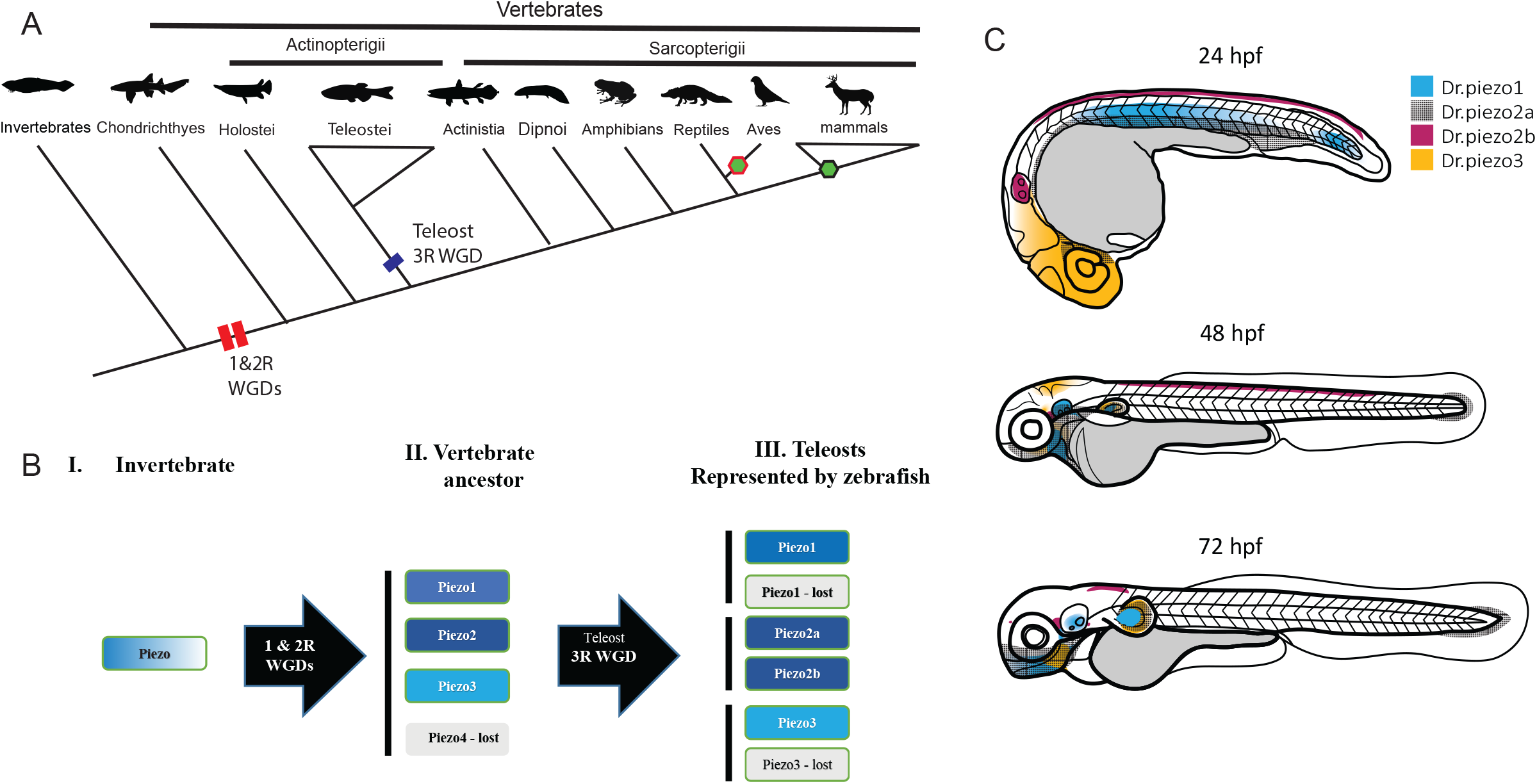
Model of vertebrate *piezo* gene evolution and their gene expression summary in zebrafish embryogenesis. **A**. The *piezo3* gene is present in most vertebrate lineages but is lost in birds and most mammals (hexagons). Red and blue bars indicated whole-genome duplications (WGDs). **B**. Based on our previous phylogenetic analysis, we proposed that there is one piezo gene in chordates. After two rounds of WGDs, four piezo genes were created in the ancestor of vertebrates. The *piezo4* gene was lost soon after the vertebrate lineage split. In the teleost lineage, the 3^rd^ WGD further created two ohnologs for *piezo1, piezo2*, and *piezo3*. However, one of piezo1 and piezo3 was lost, whereas piezo2a and piezo2b were retained in almost all teleost species. The gray colored boxes indicate gene loss. **C**. Schematic summary of *piezo* gene expressions in early zebrafish embryos (1-3dpf). The gene expression domains are color-coded.

## DISCUSSION

Mechanosensing and transduction are essential for cellular and organism physiology. The PIEZO1 and PIEZO2 channels are critical for vertebrate somatosensation, embryogenesis, and the pathogenesis of various diseases. Currently, only Piezo1 and Piezo2 genes have been reported in the literature and annotated in most vertebrate genomes (3, 4). Here, we report a new member of the vertebrate Piezo channel family, Piezo3.

WGDs play a crucial role in the Evolution of modern vertebrates and flowering plants (18, 36). It is well accepted that there were two consecutive rounds of WGD (2Rs) around the origin of vertebrates (17, 18, 20, 22). These 2Rs generated four copies of all the genes in the vertebrate ancestor, and some of these ohnologs may have been lost during Evolution. This could lead to 1-4 genes in most tetrapods. For example, there is only one hedgehog gene, while mice and humans have SHH (sonic hedgehog), IHH (Indian hedgehog), and DHH (desert hedgehog) (37). Our phylogenetic and syntenic analyses revealed that Piezo3 most likely resulted from two rounds of WGDs in the vertebrate lineage, because the Piezo3 cluster is parallel to that of Piezo1 and Piezo2. Piezo3 was retained in teleosts, amphibians, and reptiles, but lost in birds and most mammals. Only a few mammals (platypus, echidna, Tasmanian devil, armadillo, red deer, and tree shrew) still carry this gene. It would be interesting to know whether the Piezo3 has a unique function or is just retained as a redundant gene in these species. The WGD also occurred within vertebrate lineages. The teleost-specific WGD, 3R, occurred in the ancestor of modern teleosts around 300 million years ago (20, 38-41). This is the reason why most teleosts, including zebrafish, have 6-8 *Hox* gene clusters (42, 43). According to our phylogenetic and syntenic analysis, the *piezo2* gene was further duplicated into *piezo2a* and *piezo2b* in this lineage, as two paralogs that were clustered with tetrapod *PIEZO2*. Based on our results, we propose a model for the presence of the *piezo3* gene in vertebrates (**Fig. 6A**). The zebrafish piezo3 originated from the 2R, and piezo2a and piezo2b originated from the 3R. The teleost *piezo1* and *piezo3* genes were most likely duplicated, but one ortholog was lost immediately after 3R, leaving only one gene in this lineage (**Fig. 6B**).

Gene expression typically corresponds to its function during embryonic development. Both regulatory and coding regions of the ancestral *piezo* gene can be duplicated through WGDs. While coding regions generally evolve more slowly than regulatory regions due to functional constraints (44). Thus, duplicates have three fates: loss, sub-functionalization, and neo-functionalization based on the DDC model (27). Consistently, we observed some overlaps among them (**Fig. 6C**). For example, *piezo1, piezo2a*, and *piezo3* are all expressed in the pectoral fin buds, while only *piezo2a* is expressed in the caudal fin buds, suggesting a sub-functionalization. Among the four *piezo* genes, *piezo2a* has distinct expression in Rohon-Beard neurons. This could be a case of neo-functionalization, but it needs to be confirmed by examining the piezo gene expression in cephalochordates and urochordates in the future.

Since piezo3 is a new channel, further studies are needed to confirm its functions. Our present data provide an essential foundation for such inquiry. Calcium signaling in transfected human HEK293T cells suggests that zebrafish Piezo3 is a cation channel, like human PIEZO channels. It is also sensitive to mechanical stimulation and can be inhibited by streptomycin, suggesting that it has evolutionarily conserved functions. Interestingly, the *piezo3* mutant larvae were more sensitive to tapping mechanical force and showed greater displacement in our VMR assays. So, losing Piezo3 made the fish more sensitive to tapping force and more active during locomotion in the light phase, but not in the dark phase. On the other hand, our *piezo3* CRISPR mutant zebrafish are viable as adults and show no obvious morphological defects, even in regions where *piezo3* is expressed during embryogenesis, such as the eyes, pharyngeal arches, and fin buds. Thus, our results support that piezo3 is not a developmentally essential gene. One explanation is that zebrafish have four *piezo* genes, each with redundant functions, as reflected in their overlapping gene expression domains (**Figs. 3, S3-S5**). This overlap may not be limited to early embryogenesis but could also occur during later organogenesis. Further functional and anatomic analysis is needed for deciphering Piezo3’s physiological functions. From a functional perspective, the nature of being a developmental nonessential gene may underlie PIEZO3 loss in birds and mammals without disrupting normal physiology. Overall, these results suggest that Piezo3 may have a repressive role in force sensing and photopic response. Its absence in birds and most mammals may facilitate their adaptation to their environment compared to other lower vertebrates.

## MATERIALS AND METHODS

### Zebrafish strains, husbandry, and CRISPR mutation

Zebrafish were raised and maintained in accordance with AAALAC-approved standards at the Purdue animal housing facility, following protocols approved by the Purdue Animal Care and Use Committee (PACUC, #1210000750). All the zebrafish experiments were carried out in a wildtype TAB (AB/Tübingen, RRID: ZIRC_ZL1) background (45, 46). Zebrafish were maintained according to the Zebrafish Book (47), and all zebrafish embryos were staged based on the Kimmel staging guide (48).

The *piezo3* loss-of-function mutants were generated using our established CRISPR method (34). Briefly, gRNAs were designed against exons 14 and 29, and DNA templates were synthesized by IDT DNA. The gRNAs were prepared using HiScribe® T7 High Yield RNA Synthesis Kit (NEB E2040L). Then, gRNAs were co-injected with the Cas9 protein (PNA Bio, CP01) into the single-cell stage of fish embryos. The T7E1 assay was used to determine mutations before the fish embryos were raised to adulthood as F0 founders, so that their offspring, F1 fish, will be further screened using the T7E1 assay. Positive F1 adult fish will be established as founder fish lines after sequencing validation.

### Bioinformatics and phylogenetic analysis

Zebrafish Piezo channel protein sequences were first identified by searching the (RRID: SCR_006472) and Ensembl (RRID: SCR_002344) databases using human gene symbols. Any zebrafish Piezo channels with partial sequences in Ensembl were further searched for full-length sequences using BLASTp in NCBI. All the retrieved zebrafish *piezo* genes were also validated in ZFIN (RRID: SCR_002560). Orthologous and paralogous genes were retrieved from Ensembl using the comparative genomics function in GRCz11. For all the orthologous protein sequences, representative protein sequences from the major metazoan taxa were selected based on their taxonomic positions and the quality of available protein sequences. When multiple isoforms were present, the longest sequence was chosen for analysis. Piezo3 orthologs were identified by BLAST search in both NCBI and Ensembl. The final protein sequences selected for the final analysis are listed in Supplementary **Table S1**.

Phylogenetic and syntenic analyses were performed as we described previously (25, 49-52). Briefly, multiple protein sequences were aligned using the MUSCLE program with default settings (53). To identify the best evolutionary model for phylogenetic analysis, we conducted a model selection test using maximum likelihood (ML) with default parameters in MEGA (RRID: SCR_000667) (54). The models with the lowest B.I.C. (Bayesian Information Criterion) scores were considered to describe the substitution pattern the best, and JTT (Jones-Taylor-Thornton model) +G (gamma distribution) was chosen. Then, we constructed phylogenetic trees using PhyML3.1 (55). ML phylogenetic analysis was performed using JTT + G with 1000 bootstrap replicates (49, 50). The final phylogenetic trees were viewed and generated with FigTree (RRID: SCR_008515) V1.4.2 (http://tree.bio.ed.ac.uk/software/figtree). The syntenic analysis was first carried out in the Genomicus browser (version 100.01). Individual gene position was then verified in Ensembl, UCSC genome browser (RRID: SCR_011791), and the Synteny Database (56).

For the *PIEZO1P2* RNA (ENSG00000237121.1), gene expression data were retrieved from the GTEx Portal: https://www.gtexportal.org. Protein tertiary structures of zebrafish Piezo1, Piezo2a, Piezo2b, Piezo3, and human PIEZO1 and PIEZO2 were generated using ColabFold: AlphaFold2 (https://www.alphafold.ebi.ac.uk, RRID: SCR_023662) with MMseqs2 (v1.5.5) (57, 58). Full-length amino acid sequences were submitted to the ColabFold notebook using default parameters.

### Zebrafish gene cloning

Partial sequences of zebrafish *piezo* genes were cloned from a mixed cDNA pool. Briefly, About 100 embryos (1-3 days post fertilization, dpf) were dechorionated, pooled, and used for total RNA extraction using TRIzol reagent (Thermo Fisher, 15596026) according to the manufacturer’s guidance. The total RNA amount and quality were checked using a Nanodrop spectrophotometer (RRID: SCR_018042). Reverse transcriptions were performed with the SuperScript® III First-Strand Synthesis System (Thermo Fisher, 18080051) following manual instructions. Phusion® High-Fidelity DNA Polymerase master mix (New England Biolabs, M0531) or PrimeSTAR® GXL DNA Polymerase (Takara Bio, R050A) was chosen for PCR amplification. Right-sized PCR products were examined on DNA agarose gels and purified with NucleoSpin Gel and PCR Clean-up Kit (Takara Bio, 740609.250). Purified PCR products of each gene were then cloned into the pJet1.2 vector (Thermo Fisher, K1232) according to the manual instructions. Correct clones were verified for correct insertional gene orientation by endonuclease digestion and sequencing. The full length of the zebrafish *piezo3* ORF was synthesized and cloned into the pCDNA3.1 vector by Gene Script to reduce random mutations of long-range PCR. Finally, the *piezo3* ORF was subcloned into the pBac vector carrying an IRES-EGFP tag to make a plasmid construct, pBAC-*piezo3*-IRES-EGFP.

### Whole mount *in situ* hybridization, cryosection, and imaging

Riboprobe DNA templates were prepared either by PCR amplification or plasmid linearization with an endonuclease at the 5’ end of the protein-coding region in the pENTR-D or pDONR221 vector. DNA templates were purified before *in vitro* transcription. Anti-sense riboprobes were synthesized by in vitro transcription using T7 RNA polymerase (Thermo Scientific, EP0111) and DIG RNA Labeling Mix (Millipore-Sigma, 11277073910) according to the manufacturers’ manuals. All synthesized riboprobes were purified using Sigma Spin post-reaction clean-up columns (Sigma, S5059) and stored at -80°C in a freezer before use.

Whole-mount *in situ* hybridizations were performed according to our established method, with some modifications (16, 25, 26). Briefly, chorions were removed using pronase (Sigma, PRON-RO) treatment before fixation for 0.5-3 dpf fish embryos. All fish embryos were staged and then fixed with 4% PFA (paraformaldehyde) for 1-2 days at 4°C, followed by dehydration in serial methanol (25%, 50%, 75%, and 100%) with PBST (Phosphate-buffered saline solution with 0.1% tween-20). Dehydrated embryos were stored in 100% methanol at -20 °C until use. We rehydrated fish embryos using a methanol gradient in PBST (100%, 75%, 50%, and 25%) for *in situ* hybridization experiments. Embryos (12-24 hpf) were then bleached with 6% H_2_O_2_ in PBST and permeabilized with proteinase K (10 μg/ml in PBT). After permeabilization, embryos were washed with PBT and fixed in 4% PFA with 0.2% glutaraldehyde for at least 3 hrs at room temperature to inactivate proteinase K. Next, fish embryos were washed with PBST (2 × 10 minutes) to remove fixatives. Then, fish embryos were incubated in pre-hybridization solution [50% formamide, 5XSSC (0.75M NaCl, 75mM sodium citrate, pH 7.0.), 2% Roche blocking powder, 0.1% Triton-X, 50 mg/ml heparin, 1 mg/ml Torula yeast RNA, 1 mM EDTA, 0.1% CHAPS, DEPC-treated ddH_2_O] overnight at 65°C in hybridization oven with gentle shaking (40 rpm). Riboprobes (1-2 µl) were added on the second day in the morning. Then, embryos were further hybridized for 48-72 hours before washing unbound riboprobes with 2X SSC and 0.2X SSC (3 × 30 minutes for each solution). Next, fish embryos were washed with KTBT 2x for 10 minutes each (50 mM Tris-HCl, pH 7.5, 150 mM NaCl, 10 mM KCl, 1.0% Tween-20) before antigen blocking using 1-2% sheep serum (Sigma, S2263) and 2 mg/ml BSA in KTBT for at least 3 hours. Anti-digoxigenin antibody conjugated to alkaline phosphatase (Roche, 11093274910) was added to the blocking solution at a 1:5000 dilution, and embryos were incubated overnight at 4°C with gentle shaking. After 6 times 1 hr washing with KTBT and overnight KTBT incubation at 4°C with gentle shaking, the following day, color reaction was carried out in NTMT solution (100 mM Tris-HCl, pH 9.5, 50 mM MgCl2, 100 mM NaCl, 0.5% Tween-20) with 75 mg/ml NBT, 50 mg/ml BCIP, and 10% DMF (N, N-dimethylformamide). For 2-3 dpf fish embryos, 1mM levamisole was added to reduce the effect of endogenous alkaline phosphatase. Color development was carried out in the dark with gentle rocking. We closely monitor the color density of each reaction. Once the embryos developed suitable color densities, the reaction was stopped with NTMT washing. The finished samples were imaged immediately or stored in 4% PFA for later imaging. For histological analysis, post-hybridization embryos were equilibrated overnight in 15% sucrose, then in 30% sucrose with 20% gelatin, after which they were embedded in 20% gelatin for cryosectioning (10-25 μm) on a cryotome. Images were acquired using an Axiocam 305 color camera on a Zeiss Stereo Discovery V12 (RRID: SCR_027509) and an Axio Imager 2 (RRID: SCR_018876) compound microscope. Whole-mount embryos were imaged in 3% methylcellulose.

### Cell Culture, Transfection, and Calcium Signal Measurement

HEK293T (RRID: CVCL_0063) cells were cultured in DMEM (Sigma-Aldrich, #D6429) supplemented with 10% FBS (Gibco, #A3840001) in a cell culture incubator at 37 °C with 5% CO_2_. The full-length zebrafish *piezo3*, pBAC-*piezo3*-IRES-EGFP (10 μg) and pGP-jRGECO1a (1.5 μg, PRID: Addgene #61563) were co-transfected into HEK293T cells at the 50% cell confluence in the 10-cm cell culture plate (Corning, #353003) using 20μl TurboFect (Thermo Scientific, #R0533) according to the manufacturer’s manual. Transfection efficiency was assessed by measuring GFP and RFP signals around 48h after transfection. The isolated single cells of RFP+/GFP+ and RFP+/GFP-were selected using a Nikon Eclipse Ti2 inverted microscope (RRID: SCR_021068) with a polygon 1000 pattern illuminator and an ORCA-Fusion digital camera. The videos and images were captured at 488 × 488 pixels (0.5 frame/s) using a Crestoptics CICERO spinning disk confocal microscope and NIS-Elements Advanced Research (RRID: SCR_027181) under the RFP (Excitation: 568 nm; Emission: 593-668 nm) and GFP (Excitation: 488 nm; Emission: 500-550 nm) channels.

To assess the basal calcium transient level without stimulation, the RFP signal from selected cells was recorded for 3 minutes, 48h after transfection. For mechanical stimulation, a borosilicate glass capillary (1.0□mm outer diameter) was generated using a Flaming/Brown micropipette puller (Model P-1000, Sutter Instrument, RRID: SCR_016842), and the tip was blunt-ended by brief flaming. The blunt-ended capillary was then mounted on a manual micromanipulator (WPU, M3301) and used to apply transient mechanical force to selected cells. To ensure comparable responses, adjacent RFP+/GFP+ and RFP+/GFP-cells were simultaneously stimulated using a single poking event. RFP calcium signals from the paired adjacent RFP+/GFP+ and RFP+/GFP-cells were continuously recorded for 4 min with 2-second intervals following the poking event. The inhibitory effect of streptomycin was evaluated at low (172 μM) and high (1032 μM) concentrations on the calcium signal after 2 hours’ administration in cell culture dishes. Streptomycin was supplied from a Penicillin–Streptomycin stock solution, corresponding to approximately 17200□μM streptomycin (10,000□μg/mL, Gibco, #15140122). To quantify calcium signals, cytoplasmic red fluorescence was manually defined and measured using ImageJ (RRID: SCR_003070) (59). Briefly, the red channel was extracted from each frame’s RFP images and converted to 8-bit grayscale for fluorescence intensity analysis. Regions of interest (ROIs) corresponding to the cytoplasm of the paired RFP+/GFP+ and RFP+/GFP-cells were manually outlined and kept consistent throughout the analysis. For each frame, the mean pixel intensity within the ROI was calculated and used to represent the RFP signal intensity at the corresponding time point, with fluorescence intensities ranging from 0 to 255 in 8-bit grayscale images. Time-course fluorescence changes were subsequently analyzed to evaluate calcium responses following the mechanical poking events. Calcium responses for each cell were quantified as the change in fluorescence relative to baseline fluorescence (Δ F/F0), where ΔF=F-F0. F represents the RFP fluorescence intensity at a given time point, and F0 represents the baseline RFP fluorescence intensity at the beginning of the recording. To evaluate the overall calcium response during the recording period, the total area under the curve (AUC) of the Δ F/F0 traces was calculated, including both positive and negative deflections relative to the ΔF/F0=0 baseline. The student t-test was performed in GraphPad Prism 9.0 (RRID: SCR_000306) for statistical analysis.

### Bone and cartilage staining

Fish were euthanized with an overdose of MS-222 and fixed in a freshly prepared solution containing 5% formalin, 5% Triton X-100, and 1% KOH. Specimens were rocked at 42 °C for 48h until pigmentation was removed. For adult fish bone staining, after a brief rinse with deionized water, calcified bone was stained with 0.02% Alizarin Red in 1% KOH for 2–3 days with gentle rocking (60). During staining, scales were removed under a binocular microscope using fine tweezers to facilitate visualization of the internal skeleton. For larvae and juvenile cartilage and bone staining, a modified double staining protocol was used (61). A double-stain solution was prepared by mixing 10 ml of Alizarin red solution (0.5% Alizarin Red in H2O) and 1ml of Alcian blue solution (0.02% Alcian blue, 200 mM MgCl2, 70% Ethanol). Specimens were incubated in the double-stain solution with gentle rocking for 1-2 days, depending on the staining stage. For both bone and cartilage staining, specimens were then washed in a clearing solution (20% Tween-20, 1% KOH) at 42 °C for 6 h until background coloration was eliminated. Samples were subsequently transferred through a graded glycerol series (25%, 50%, 75%) and stored in 100% glycerol for immediate imaging or long-term preservation. Images were acquired using an Axiocam 305 color camera on a Zeiss Stereo Discovery V12 microscope.

### Visual-Motor Response (VMR) and Tapping Assay

At 6 dpf, wildtype and homozygous mutant zebrafish larvae were transferred to Whatman UNIPlate square 96-well plates (VWR, Radnor, PA) along with their respective E3 treatment medium, with one larva per well and 24 larvae per treatment group. The plates were placed in lightproof boxes inside the incubator for overnight dark habituation. At 7 dpf, they were transferred to the Zebrabox (Viewpoint Life Sciences, Montréal, QC, Canada) to measure the visual-motor response (VMR) displayed by the larvae. The total irradiance of the LED stimulus across the visible spectrum (380–780 nm) was 310.45 µW/cm^2 (62). Larval movement was recorded in tracking mode, which binned activity every second. The data collection protocol consists of a 30-minute dark period, a 1-hour light period, and a 5-minute dark period. All VMR experiments were conducted between 9 a.m. and 6 p.m. to minimize circadian rhythm effects on vision (63, 64). The VMR data were first normalized to remove confounding factors, including light-intensity variation across the 96-plate and batch effects across biological replications using an established procedure (65). We first analyzed the displacement during the first second after light onset and offset, a typical approach for detecting visual startle responses in larvae, using a linear mixed-effects model with genotype as the fixed effect and biological replicate as the random effect (66). The results showed no significant effect of the genotype on larval displacement (on: β = -0.01341, p = 0.606; off: β = 0.07686, p = 0.0974). We also noticed a substantial difference in locomotor activities during the 1-hour light period. We analyzed this difference using the total displacement during the 1-hour period in a linear mixed-effects model, with genotype as the fixed effect and biological replicate as the random effect. The results showed a non-significant biological replicate effect and a significant increase in piezo3 mutant displacement (β= 63.2, p = 0.0065). Since the biological replicate effect was non-significant, we combined the total displacement values for each genotype from all replicates and visualized the combined data in a boxplot (Figure 9D).

A tapping assay was conducted to evaluate the total distance traveled by larvae in response to a mechanosensory stimulus, i.e., a physical tap. These larvae were transferred to 96-well plates and placed in the Zebrabox as described in the VMR assay section. To induce mechanosensory stimulation, a metal rod was manually tapped against the side of the 96-well plate under constant ambient light. Each experiment consists of 12 taps, with a 10-second interval between taps. Larval responses were recorded by the Zebrabox in tracking mode. Each plate was tested twice. During the recording, the larvae were exposed to constant ambient light. The tapping assay data were analyzed in two steps: 1) filtering and 2) hypothesis testing. First, the total distance data were filtered to remove unsuccessful taps. A tap was considered unsuccessful if the average total distance traveled at that tap was less than two standard deviations above the average total distance traveled during periods between taps. The average total distance during the in-between periods was calculated across all genotypes and all in-between time intervals. Second, the filtered data from all taps and plates were pooled within each genotype. The Wilcoxon rank-sum test was used to test the difference in average total distance traveled between genotypes. The resulting p-values were adjusted using Holm’s method to correct for multiple comparisons (67).

## ACKNOWLEDGMENT

The research was supported by the National Institute of General Medical Sciences of the National Institutes of Health (R35GM124913) to G.Z. The content is solely the responsibility of the authors and does not necessarily represent the official views of the funding agents. The authors also thank the Hayward Foundation for its support of our research.

## AUTHOR CONTRIBUTIONS

**Z.D**., **D.W**., **B.W**., **J.A.N**., and **Y.F.L**.: Formal analysis, Investigation, Methodology, Writing-review & editing. **G.Z**.: Conceptualization, Formal analysis, Funding acquisition, Investigation, Supervision, Writing-original draft, Writing-review & editing.

