## Supplemental figures for "Piezo3 is a novel mechanosensitive Piezo ion channel in vertebrates"

Dong et. al.

#### Supplementary files

**Figure S1. An extended majority-rule consensus tree for the Bayesian phylogenetic analysis of animal PIEZO channels.** Some representative vertebrate species (human, mouse, zebrafish, spotted gar, shark) and a few invertebrates were analyzed. Numbers at each node indicate posterior probability (pp) values based on twenty million runs. Branch lengths are proportional to the means of the pp densities for their expected replacements per site. The ML phylogenetic tree (**Fig. 1**) was generally consistent with this BP phylogeny: all invertebrates have one PIEZO, while there are three PIEZO (PIEZO1-PIEZO3), which are color-coded and indicated with vertical black lines. There are extra Piezo2 proteins in teleosts. The invertebrates form the outgroups of vertebrates.

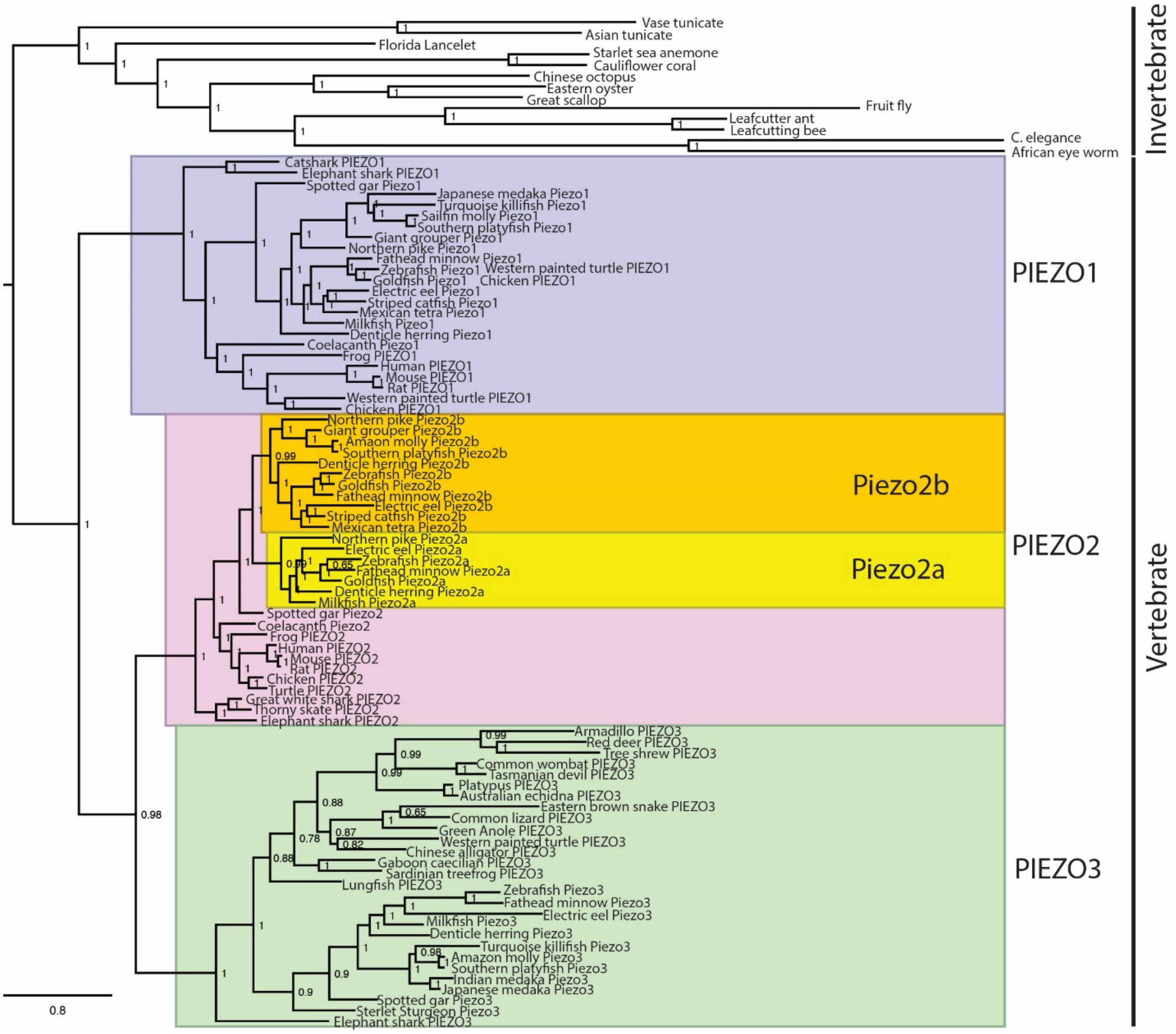

**Revealing Piezo3 as a new mechanosensitive Piezo ion channel in vertebrates**

Dong et. al.

**Figure S2. Tissue distribution of the human *PIEZO1P2* transcript expression.** The data was exported from the GTEx Portal database using default parameters.

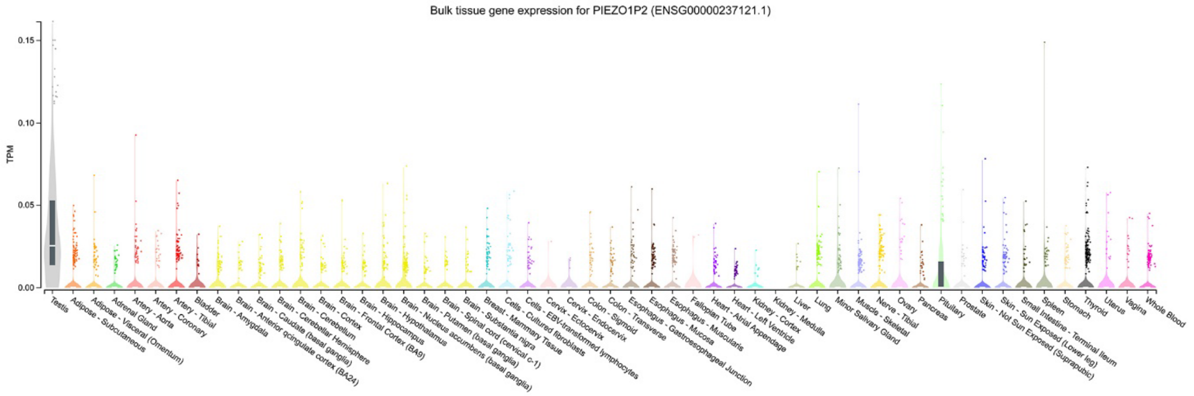


**Figure S3. Zebrafish *piezo1* gene expression during zebrafish embryogenesis.** Whole-mount in situ hybridization of zebrafish embryos at bud stage (**A**-**C**), 5 somite stage (**D**-**F**), 15 somite stage (**G**-**I**), 21 somite stage (**J**-**K**), 24hpf (**L**-**M**), 48hpf (**N**-**O**), and 72 hpf (P-Q). The anterior is to the left in all the whole-mount images, and the dorsal is to the top in all transverse sections (**L**, **N**, **P**). The gene expression of *piezo1* is viewed laterally (**A**, **D**, **G**, **J**, **L**, **N**, **P**). Dorsal view of gene expression of *piezo1* (**B**, **E**, **H**, **K**, **M**, **O**, **Q**). Ventral view of gene expression of *piezo1* (**C**, **F**, **I**). The white dashed lines indicate the proximate positions of the sections shown in the insert (**L'**) of the same panel (**L**). Magnified pectoral fin buds are highlighted in the inserts (**N'**, **P'**) in the corresponding panels (**N**, **P**). Scale bars were added to the left of the bottom row. 250 µm for whole-mount images. *e*, eye; *en*, endoderm; *fb*, forebrain; *hb*, hindbrain; *mb*, midbrain; *n*, notochord; *nt*, neural tube; *op*, optic vesicles; *ov*, otic vesicles; *pa*, pharyngeal arches; *pf*, pectoral fin bud; *sp*, subpallium; *tb*, tail bud; *vm*, ventral mesoderm of somite.

**Revealing Piezo3 as a new mechanosensitive Piezo ion channel in vertebrates**

Dong et. al.

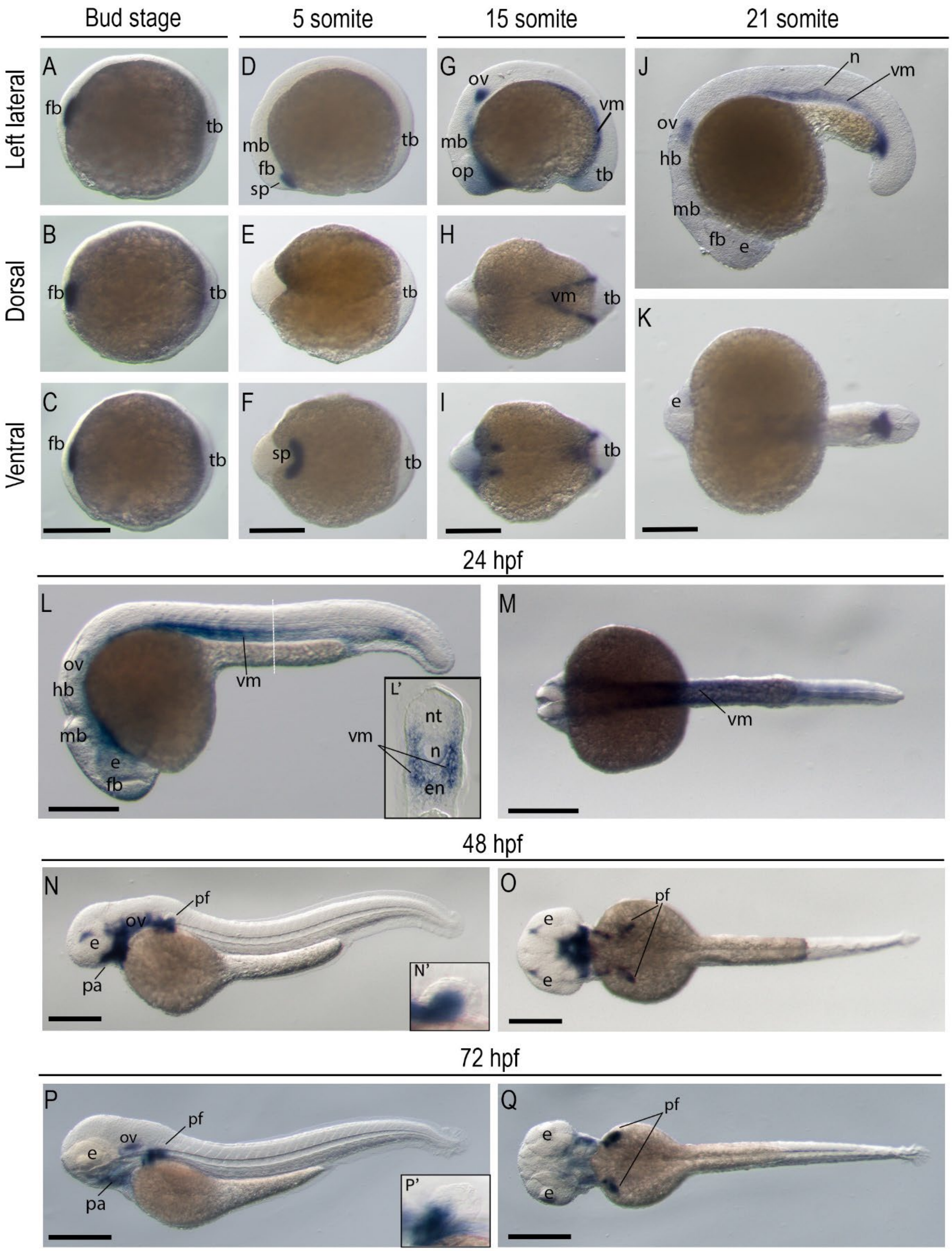

**Revealing Piezo3 as a new mechanosensitive Piezo ion channel in vertebrates**

Dong et. al.

**Figure S4. Zebrafish *piezo2a* gene expression during zebrafish embryogenesis.** Whole-mount in situ hybridization of zebrafish embryos at bud stage (**A**-**C**), 5 somite stage (**D**-**F**), 15 somite stage (**G**-**I**), 21 somite stage (**J**-**K**), 24hpf (**L**-**M**), 48hpf (**N**-**O**), and 72 hpf (**P**-**Q**). The anterior is to the left in all the whole-mount images, and the dorsal is to the top in all transverse sections (**L**, **N**, **P**). The gene expression of *piezo2a* is viewed laterally (**A**, **D**, **G**, **J**, **L**, **N**, **P**, **R**). Dorsal view of gene expression of *piezo2a* (**B**, **E**, **H**, **K**, **M**, **O**, **Q**, **S**). Ventral view of gene expression of *piezo2a* (**C**, **F**, **I**). The white dashed lines indicate the proximate positions of the sections shown in the insert (**L'**, **M'**) of the same panels (**L**, **M**). Magnified pectoral and caudal fin buds are highlighted in the inserts (**N**'-**S'**) in the corresponding panels (**N**-**S**). Scale bars were added to the left of the bottom row. 250 µm for whole-mount images. *cf*, caudal fin bud; *e*, eye; *en*, endoderm; *fb*, forebrain; *hb*, hindbrain; *hm*, head mesenchymal; *mb*, midbrain; *n*, notochord; *nt*, neural tube; *op*, optic vesicles; *ov*, otic vesicles; *pa*, pharyngeal arches; *pf*, pectoral fin bud; *tb*, tail bud; *vm*, ventral mesoderm of somite.

**Revealing Piezo3 as a new mechanosensitive Piezo ion channel in vertebrates**

Dong et. al.

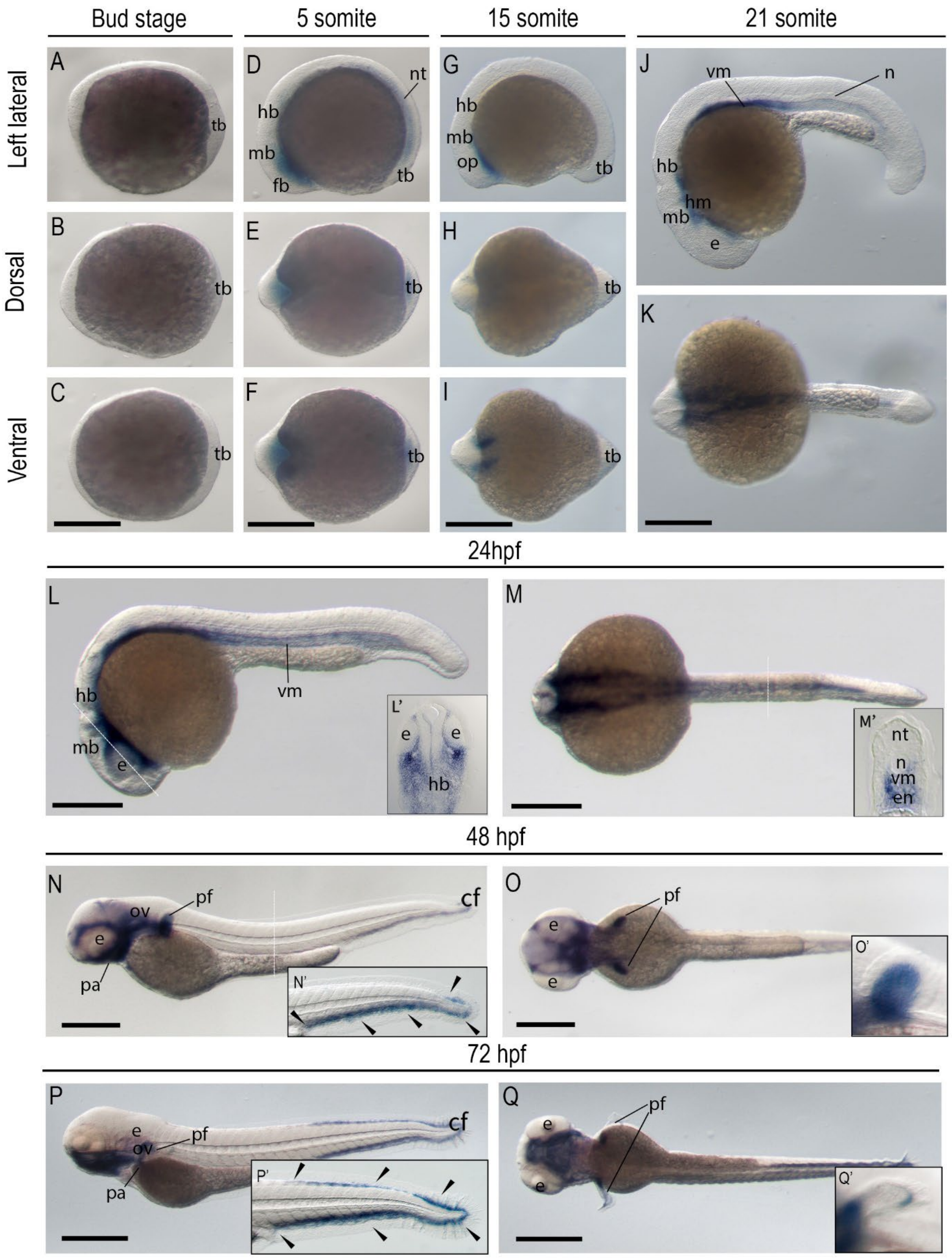

**Revealing Piezo3 as a new mechanosensitive Piezo ion channel in vertebrates**

Dong et. al.

**Figure S5. Zebrafish *piezo2b* gene expression during zebrafish embryogenesis.** Whole-mount in situ hybridization of zebrafish embryos at bud stage (**A**-**C**), 5 somite stage (**D**-**F**), 15 somite stage (**G**-**I**), 21 somite stage (**J**-**K**), 24hpf (**L**-**M**), 48hpf (**N**-**O**), and 72 hpf (**P**-**Q**). The anterior is to the left in all the whole-mount images, and the dorsal is to the top in all transverse sections (**L**, **M**). The gene expression of *piezo2b* is viewed laterally (**A**, **D**, **G**, **J**, **L**, **N**, **P**). Dorsal view of gene expression of *piezo2b* (**B**, **E**, **H**, **K**, **M**, **O**, **Q**). Ventral view of gene expression of *piezo2b* (**C**, **F**, **I**). The white dashed lines indicate the proximate positions of the sections shown in the insert of the same panel (**L-L'**, **M-M''**). Scale bars were added to the left of the bottom row. 250 μm for whole-mount images. *e*, eye; *fb*, forebrain; *hb*, hindbrain; *mb*, midbrain; *nt*, notochord; *nt*, neural tube; *op*, optic vesicles; *ov*, otic vesicles; *pa*, pharyngeal arches; *rhb*, Rohon–Beard neurons; *tb*, tail bud; *tg*, trigeminal ganglia.

### Revealing Piezo3 as a new mechanosensitive Piezo ion channel in vertebrates

Dong et. al.

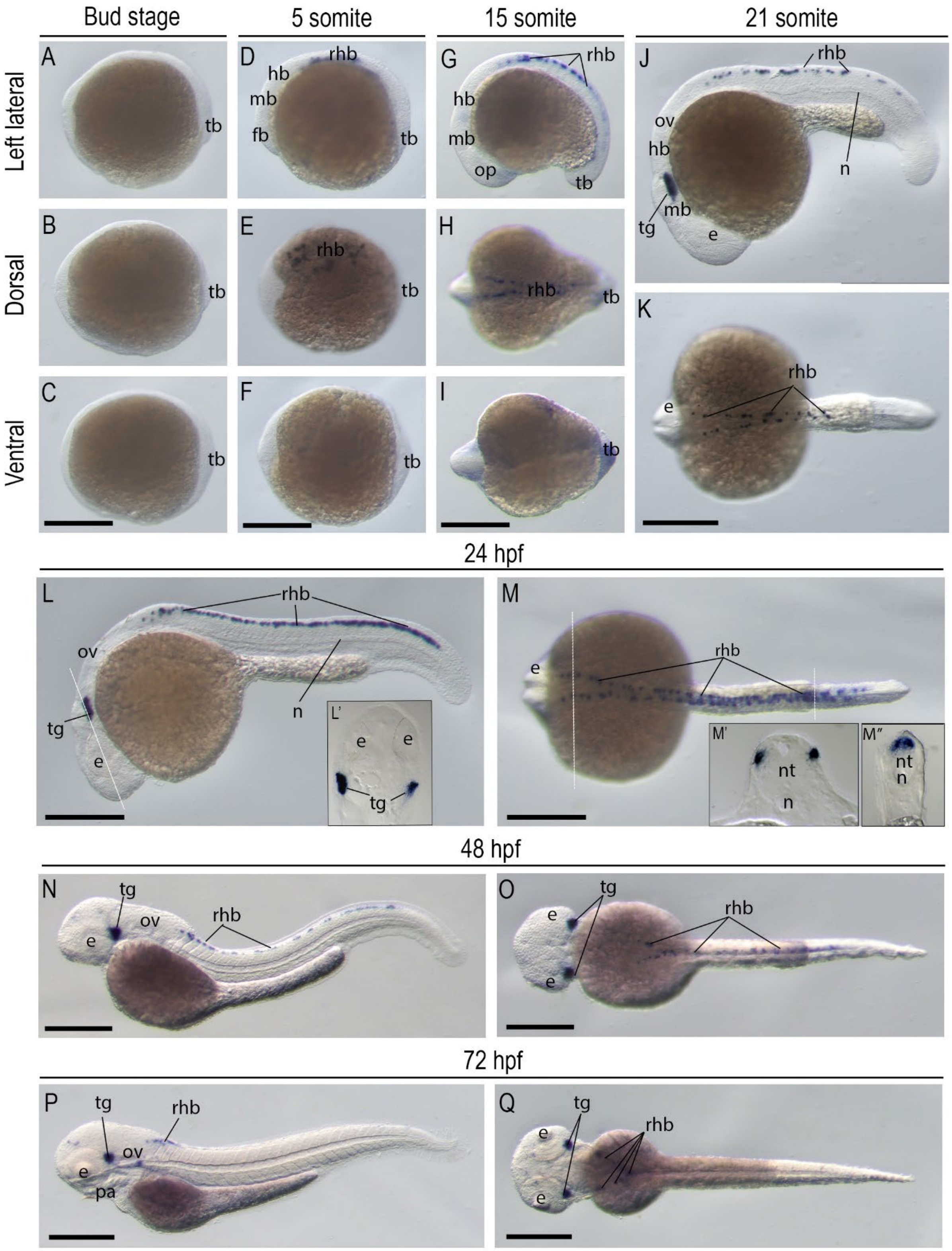

**Revealing Piezo3 as a new mechanosensitive Piezo ion channel in vertebrates**

Dong et. al.

**Figure S6. Conserved structural architecture of vertebrate PIEZO proteins revealed by AlphaFold2 prediction.** Predicted protein structure of PIEZO channel subunit from zebrafish and human using AlphaFold2. **A.** Dr.Piezo3 (this study), **B.** Hs.PIEZO1, **C.** Hs.PIEZO2, **D.** Dr.Piezo1, **E.** Dr.Piezo2a, and **F.** Dr.Piezo2b. Structural domains are annotated based on previously resolved human PIEZO1 structures and domain definitions (64, 65). *Anchor*, anchor domain is indicated in purple, linking the peripheral blades to the central pore module. *Beam*, intracellular beam domain (bright green). *CED*, central extracellular domain (light green). *CTD*, C-terminal domain (brown). *IH*, inner helix domain (red). *OH*, outer helix (blue). *THU*, transmembrane helical units (cyan), which form the propeller-like blades.

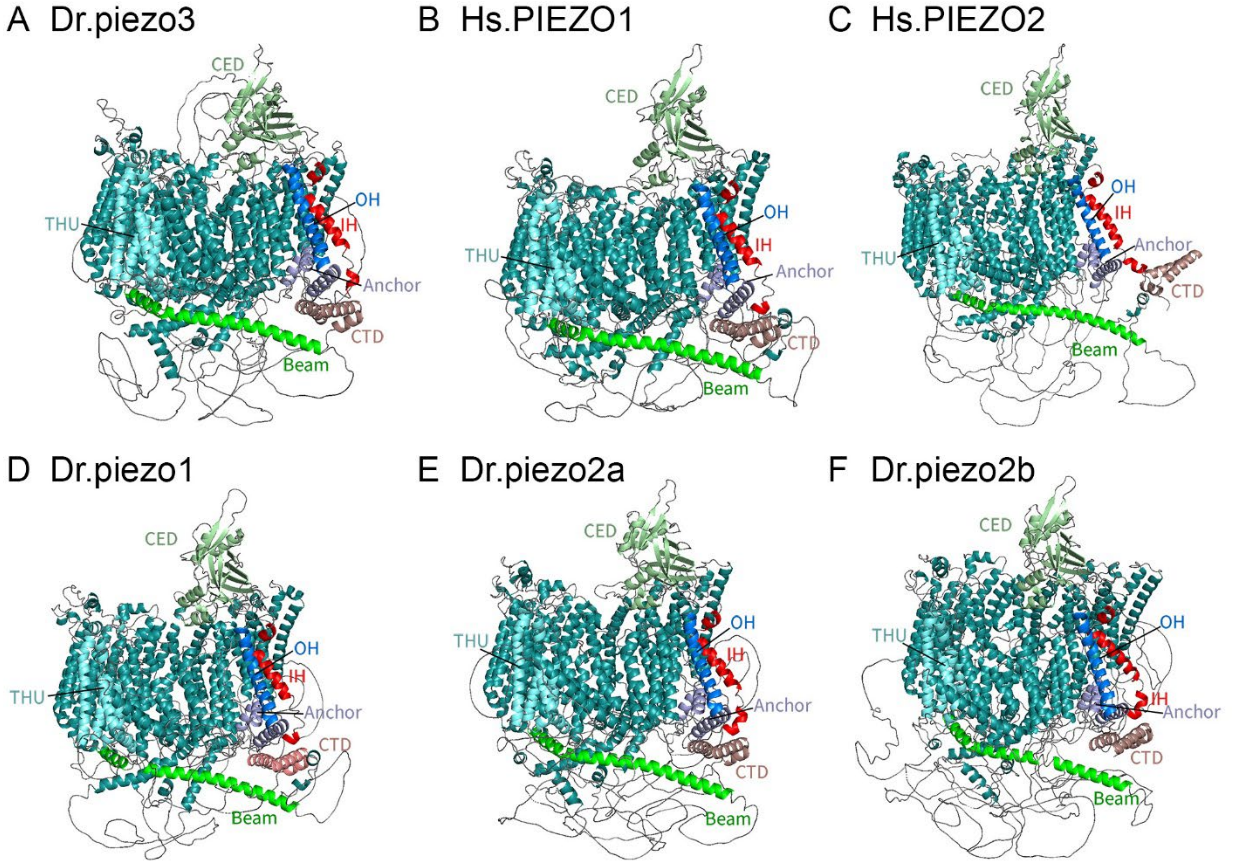
